# Native Adiponectin Plays a Role in the Adipocyte Mediated Conversion of Fibroblasts to Myofibroblasts

**DOI:** 10.1101/2022.12.23.521683

**Authors:** Mariam El-Hattab, Noah Sinclair, Jesse Liszewski, Michael Schrodt, Jacob Herrmann, Aloysius J. Klingelhutz, Edward A. Sander, James A. Ankrum

## Abstract

Adipocytes regulate tissues through production of adipokines that can act both locally and systemically. Adipocytes also have been found to play a critical role in regulating the healing process. To better understand this role, we developed a 3D human adipocyte spheroid system that has an adipokine profile similar to *in vivo* adipose tissues. Previously, we found that conditioned medium from these spheroids induces human dermal fibroblast conversion into highly contractile, collagen producing myofibroblasts through a transforming growth factor beta-1 (TGF-β1) independent pathway. Here, we sought to identify how mature adipocytes signal to dermal fibroblasts through adipokines to induce myofibroblast conversion. By using molecular weight fractionation, heat inactivation, and lipid depletion, we determined mature adipocytes secrete a factor that is 30-100 kDa, heat labile, and lipid associated that induces myofibroblast conversion. We also show that depletion of the adipokine adiponectin, which fits those physiochemical parameters, eliminates the ability of adipocyte conditioned media to induce fibroblast to myofibroblast conversion. Interestingly, native adiponectin secreted by cultured adipocytes consistently elicited a stronger level of *α*-SMA expression than exogenously added adiponectin. Thus, adiponectin secreted by mature adipocytes induces fibroblast to myofibroblast conversion and may lead to a phenotype of myofibroblasts distinct from TGF-β1 induced myofibroblasts.

## Introduction

Fibroblasts are viewed as central players in wound healing. After injury, fibroblasts, in response to biochemical factors, such as transforming growth factor beta-1 (TGF-β1), and mechanical factors such as tensile forces in the extracellular matrix (ECM), differentiate into a highly-contractile myofibroblast phenotype that rapidly remodels and repairs the tissue^1^. Normally, myofibroblasts disappear at the conclusion of healing. However, chronically activated myofibroblasts persist at the wound site and are associated with excessive collagen production and tissue contracture. Consequently, myofibroblasts remain a common target of therapies designed to prevent scar formation ^2^.

A growing body of evidence indicates that adipocytes also contribute to tissue repair and restoration both directly and indirectly through interactions with other resident cells in the wound, including fibroblasts ^3-5^. Lineage tracing in mice has revealed that adipocytes quickly repopulate the wound site, secrete growth factors, and deposit ECM proteins, all of which influences fibroblast activity and healing outcomes^5,6^. In fact, inhibition of adipogenesis was found to significantly decrease fibroblast density and activity in healing wounds, which suggests that interactions between adipocytes and fibroblasts are an essential component of the healing proccess^5^. However, the mechanisms by which adipocytes influence fibroblast/myofibroblasts during ECM remodeling remain to be elucidated.

In order to identify these mechanisms, we developed a 3D human adipocyte spheroid culture system and investigated the influence of adipocyte conditioned medium (ACM) on fibroblast behavior. We found that secreted factors contained within ACM promoted the conversion of human dermal fibroblasts to *α*-smooth muscle actin (*α*-SMA) positive, contractile, collagen producing myofibroblasts^7^. Furthermore, we determined that this conversion, which did not happen in the presence of pre-adipocyte conditioned media, occurred though a TGF-β1 independent mechanism, as small molecule inhibition of the TGF-β1 receptor did not significantly diminish fibrin gel compaction in adipocyte conditioned medium (ACM) treated samples. Since this conversion does not appear to be TGF-β1 dependent, the resultant myofibroblasts are likely distinct in their tissue remodeling potential. However, the identity of the adipocyte secreted factor responsible for conversion remains unknown.

Here, we report on our progress in identifying the factor responsible for adipocyte mediated myofibroblast conversion with a specific emphasis on the role that the adipokine adiponectin could be playing. Since the behavior of myofibroblasts are determined in part by the factors that induce their differentiation, uncovering the mechanisms by which adipocyte secreted factors induce myofibroblast conversion is a critical first step to determining if adipocytes generate myofibroblasts that are scar forming versus tissue regenerating.

## Results

### Conditioned medium from primary adipose-derived cells induces myofibroblast conversion

We reported previously on the myofibroblast inducing potential of adipocyte conditioned media derived from an immortalized human pre-adipocyte cell line (NPAD)^7^. To determine if this potential is also present in spheroids formed from primary human cells, we collected conditioned media from primary cells and quantified myofibroblast conversion similar to our previous studies^7-9^. Adipose mesenchymal stromal cells isolated from the stromal vascular fraction of digested donor adipose tissue were differentiated in spheroid cultures into either pre-adipocyte or adipocytes. After 10 days of differentiation, conditioned media was collected and assessed for its ability to induce myofibroblast conversion. We found adipocyte conditioned media (ACM) derived from both the immortalized NPAD-derived and the primary SVF-derived cultures significantly induced *α*-SMA levels above the control and similar to the positive control (Fig. 1). Consistent with our prior studies, pre-adipocyte conditioned media (PCM) did not increase *α*-SMA expression (Fig 1). Thus, primary SVF-derived human adipocytes also induce myofibroblast conversion, a finding that gives us confidence that this signaling mechanism is a general feature of our 3D spheroid culture system.

**Figure 1.**
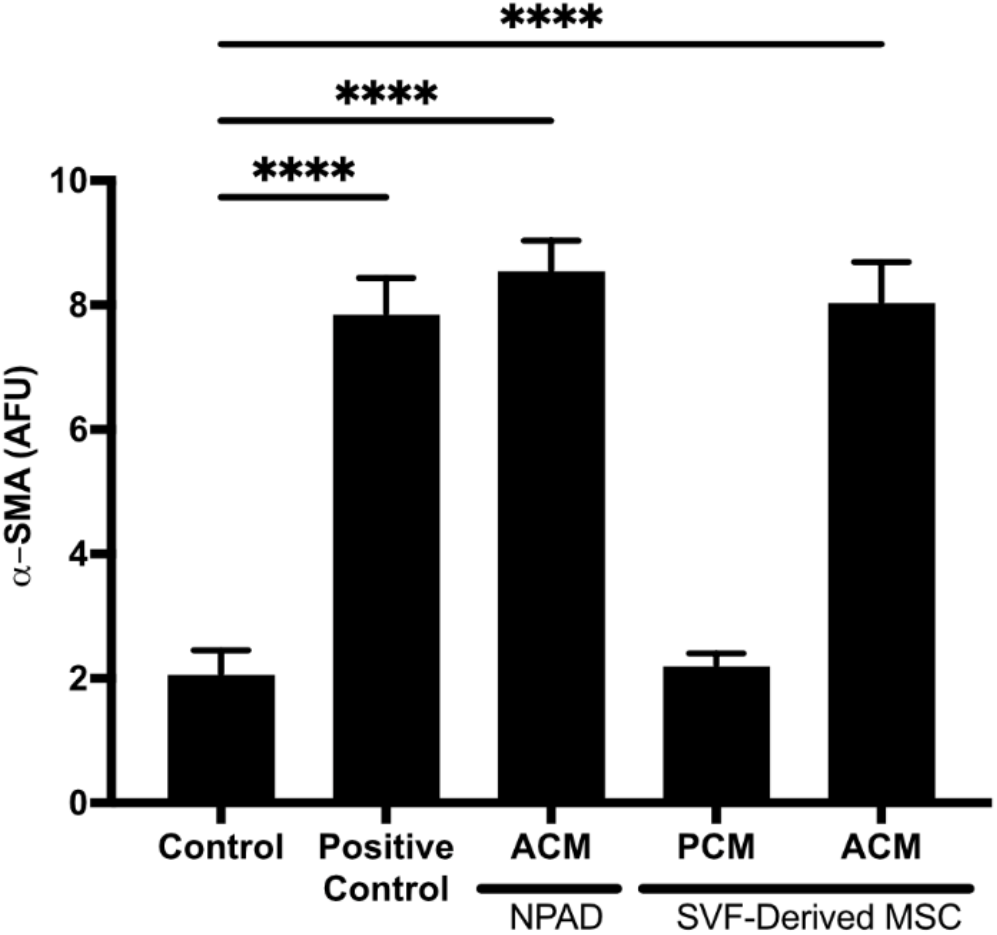
ACM from SVF-derived cultures induces myofibroblast conversion similarly to immortalized NPAD-derived cultures. ACM and PCM from SVF-derived MSC cultures were assessed for their ability to induce human fibroblast expression of *α*-SMA. ACM collectes from immotalized NPAD-derived cultures and positive control media containing TGF-β1 and ascorbic acid were used as controls. *α*-SMA expression was measured using a fluorescent plate reader after immunolabeling. One-way ANOVA with Dunnett’s multiple comparison relative to control ****p<0.0001, Data is mean ± SD, N = 3.

### ACM and PCM differ substantially in adipokine profile

Having established that ACM from primary cultures induces myofibroblast conversion while PCM does not, we investigated what factor(s) are present in ACM but not PCM that could potentially be responsible. We began by comparing the adipokine content of human SVF-derived ACM and PCM. We found that 4 of the 6 adipokines analyzed differed significantly and by orders of magnitude between ACM and PCM (Fig. 2). ACM had levels of adiponectin, adipsin, retinol binding protein-4 (RBP4), and interferon gamma (IFN-*γ*) 10-100 times higher than PCM. In contrast, PCM had slightly higher levels of interleukin 8 (IL-8) and interleukin 6 (IL-6), but the experiment was not powered enough to declare the differences in IL-8 and IL-6 statistically significant (p = 0.078 and p =0.054, respectively). Thus, elevated adiponectin, adipsin, RBP4, and IFN-*γ* are all candidate adipokines that could be playing a role in ACM-mediated induction of myofibroblast conversion.

**Figure 2.**
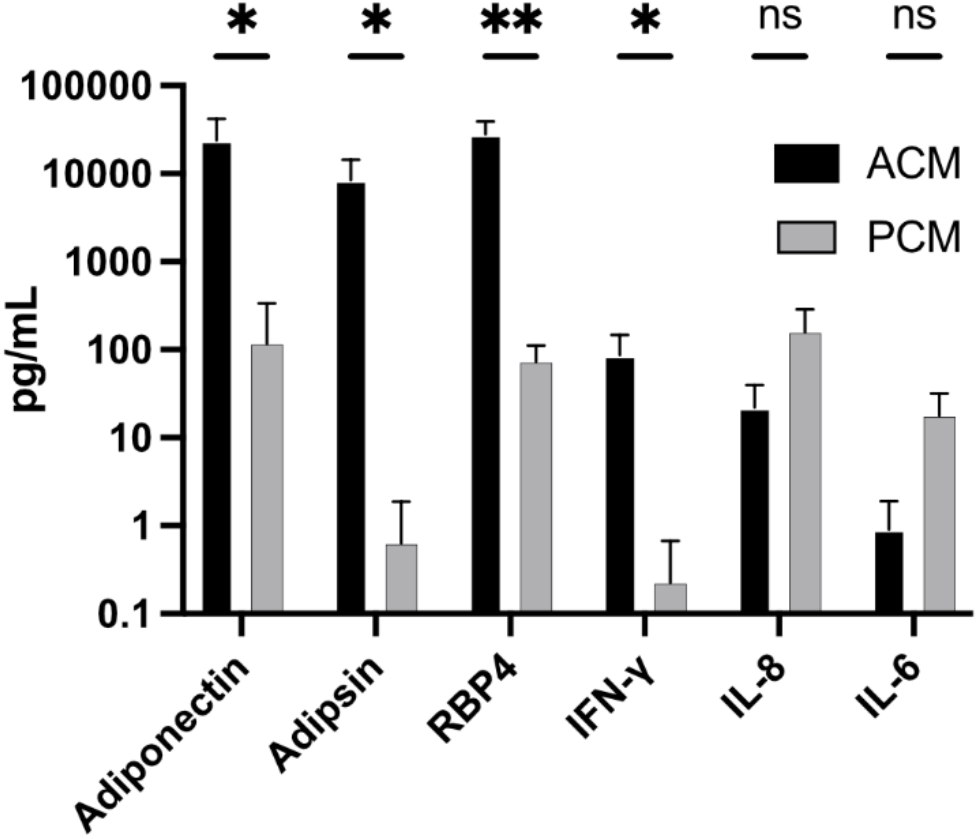
The adipokine profile between ACM and PCM differs significantly. Adipokine profile of ACM and PCM as measured using a bead-based adipokine array. Each cytokine was compared using unpaired t-tests, *p<0.05, **p<0.005. Data is mean ± SD, N=4.

### Fractionation of ACM reveals factor physiochemical properties consistent with adiponectin

Having found substantial differences between the adipokine profiles of PCM and ACM, we next performed a physiochemical characterization of conditioned media to help narrow candidate factors responsible for myofibroblast conversion. Since the adipokines elevated in ACM have different molecular weights (adiponectin – monomeric 30kDa, trimeric 90kda; adipsin – 28 kDa; RBP4 – 21 kDa; IFN-*γ* – 17 kDa, IL-6 – 21kDa; IL-8 – 8.4 kDa) we fractionated the ACM using molecular weight cutoff (MWCO) filters in the ranges of <5 kDa, 5-10 kDa, 10-30 kDa, 30-100 kDa, and >100 kDa and then repeated the *α*-SMA assay. We found that after fractionation, the ability of ACM to induce *α*-SMA expression was lost in all fractions except the 30-100 kDa fraction (Fig. 3A). Thus, we conclude that the factor could be adiponectin, but that it is not adipsin, RBP4, IFN-*γ*, IL-6, or IL-8.

**Figure 3.**
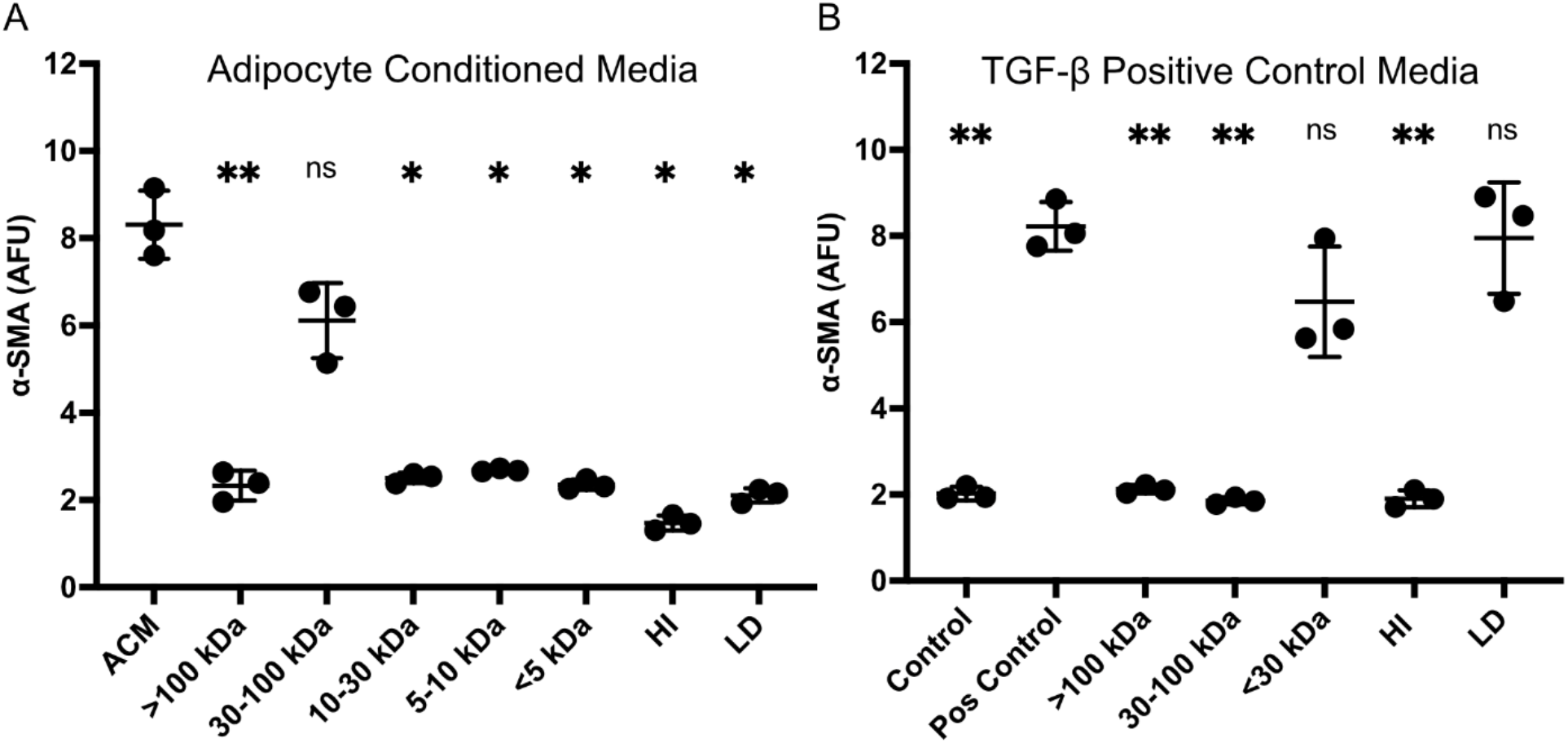
Fractionation of Adipocyte Conditioned Media reveals the inducing factor is between 30-100 kDa, heat labile, and lipid associated. A) ACM fractionation shows a molecule(s) in the 30-100 kDa fraction involved in fibroblast conversion to myofibroblasts. HI and LD both removed this effect. B) TGF-β1 Positive Control fractionation shows that the TGF-*β*1 was retained in the <30 kDa and LD fractions. Myofibroblast conversion measured via fluorescently labeled *α*-SMA. Data is mean ± SD, N=3. One-way ANOVA conducted with Dunnett’s multiple comparisons test. *p<0.05, **p<0.005.

To determine if the factor is a protein, we heat inactivated (HI) the ACM and found that doing so completely eliminated the ability of ACM to induce *α*-SMA expression (Fig. 3A). Finally, to determine if the factor is lipid associated, we depleted lipids (LD) and lipid associated factors using Cleanascite and found the *α*-SMA inducing activity of ACM was also eliminated (Fig. 3A). When fractionation, heat inactivation, or lipid depletion was performed on TGF-β1 containing positive control media, only the <30kDa fraction and lipid depleted ACM retained *α*-SMA inducing activity, findings completely expected and consistent with the physiochemical properties of TGF-β1 (the active form of TGF-β1 is a 25 kDa dimer). Thus, the factor in ACM responsible for inducing myofibroblast conversion shows distinct physiochemical characteristics compared to TGB-β1 containing positive control media, which is an additional confirmation of our earlier study^7^.

The observation that both lipid depletion and heat inactivation eliminated myofibroblast conversion suggests that both a lipid and a heat-labile protein could be involved. To test for this possibility, we combined ACM HI, which should retain the lipids, and ACM LD, which should retain the proteins, and tested for myofibroblast conversion. We again found ACM HI and ACM LD alone eliminated myofibroblast conversion but combining ACM HI and ACM LD also did not restore any measurable increase in *α*-SMA induction (Fig. S1). While this experiment does not eliminate the possibility that multiple factors are involved, it does raise the possibility that the protein itself could also be lipid associated, as it appears to be removed/deactivated during lipid depletion.

**Figure S1.**
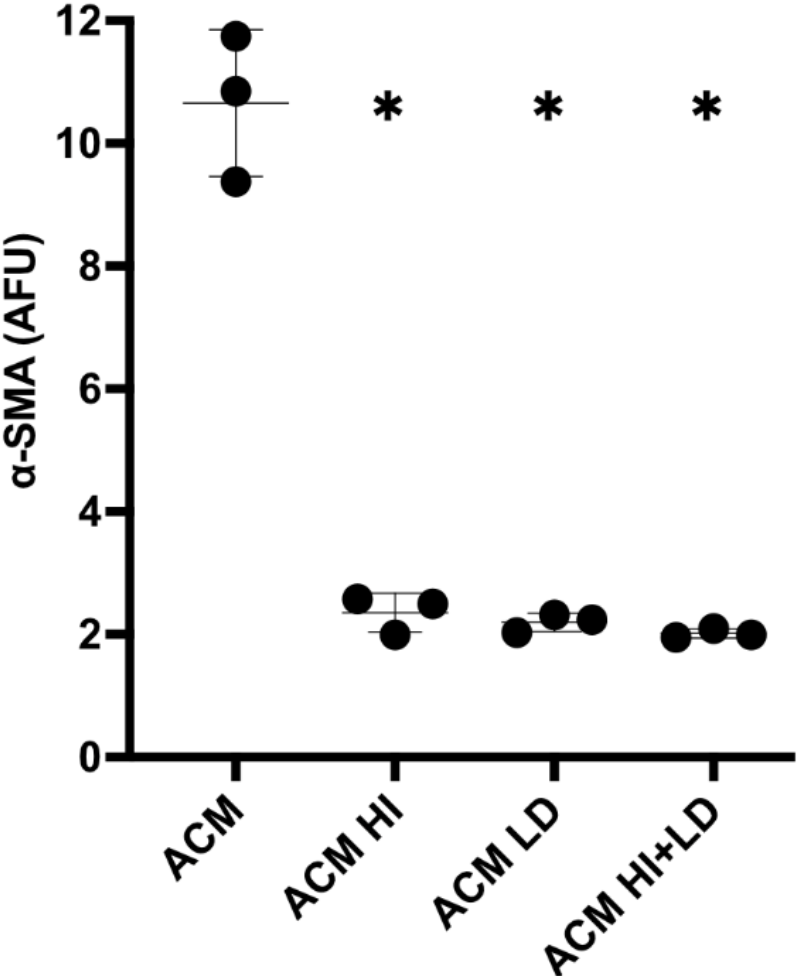
ACM factor responsible for myofibroblast induction is both heat labile and lipid associated. ACM was heat inactivated (ACM HI) or lipid depleted (ACM LD) and then applied to fibroblasts. To see if two different factors, one heat labile and one lipid associated were responsible, the HI and LD medias were combined in a 1:1 ratio and applied to fibroblasts (ACM HI+LD). Myofibroblast conversion measured via fluorescent immunolabeling of *α*-SMA. Data is mean ± SD, N=3. One-way ANOVA conducted with Dunnett’s multiple comparisons test. *p<0.05.

### High concentrations of exogenously added adiponectin can induce myofibroblast conversion

Of the four substantially elevated adipokines we identified in ACM compared to PCM, only adiponectin conforms to the 30 – 100 kDa range where myofibroblast conversion was observed. Adiponectin is a complex molecule with a monomeric molecular weight of approximately 30 kDa. It has three secreted oligomeric forms: a low molecular weight (LMW) trimer that is 67 kDa, a middle molecular weight (MMW) hexamer that is 140 kDa, and a high molecular weight (HMW) multimer that is at least 300 kDa monomers^10,11^. A bioactive, proteolytically degraded globular form that consists of the trimeric head domain minus the stalk also exists natively^12^. To add further complexity, adiponectin also can bind anionic phospholipids^13^. Based on the physiochemical properties of the unknown factor in ACM, we sought to determine if adiponectin could be the factor in ACM responsible for myofibroblast conversion by performing a dose response experiment using exogenously added human adiponectin (ex-Adip). We found ex-Adip at concentrations of 25 ng/ml and above induced significant myofibroblast conversion in a dose-dependent manner (Fig. 4A). Fitting a dose response curve to the data revealed a half maximal effective concentration (EC_50_) of 37 ng/mL^14^. Although these data indicate that ex-Adip can induce myofibroblast conversion, the calculated EC_50_ value for ex-Adip is 2-4 times higher than the concentration of native adiponectin measured in ACM.

**Figure 4.**
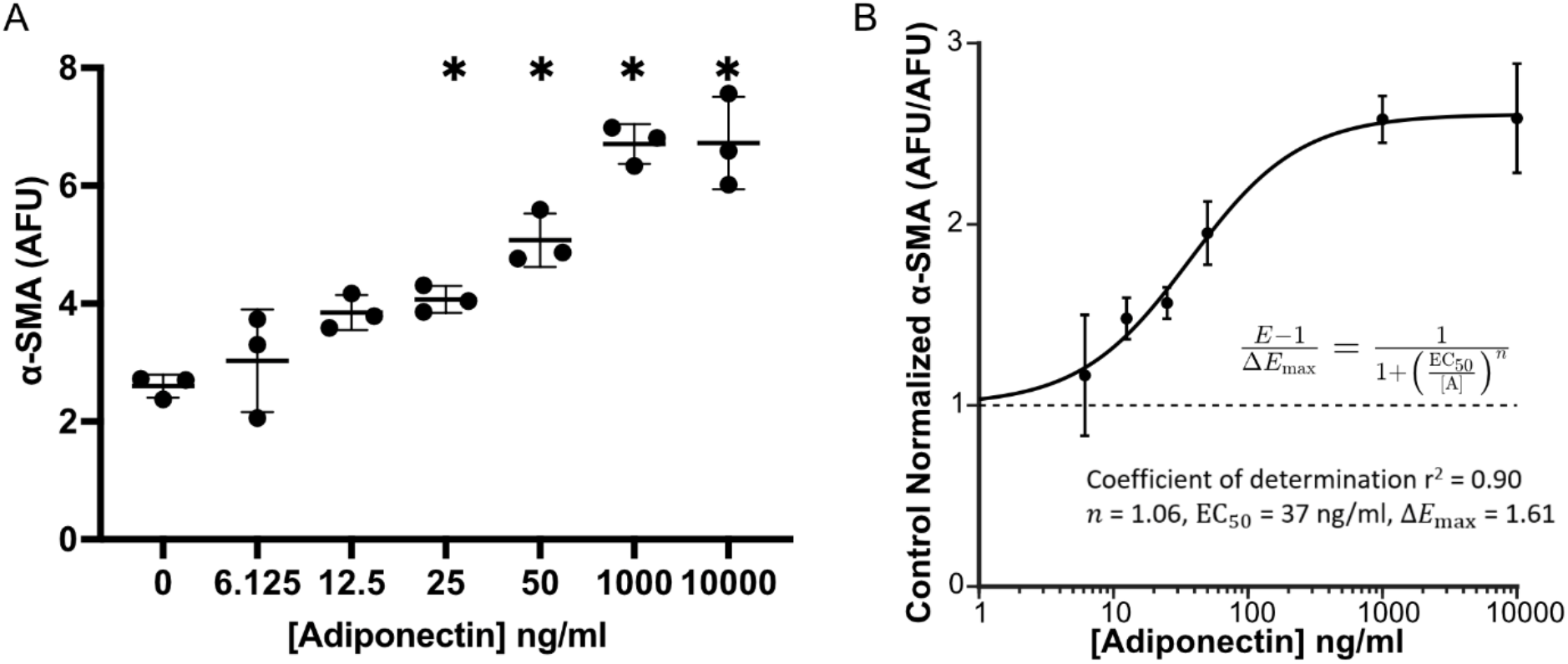
Human adiponectin promotes expression of *α*-SMA from fibroblasts in a dose-dependent manner. A) Myofibroblast conversion measured via fluorescent immunolabeling *α*-SMA in fibroblasts 48 hours after treatment with increasing concentrations of human adiponectin. B) To fit a dose response curve, control-normalized *α*-SMA expression (*E*) in response to each adiponectin concentration ([A]) was computed as the ratio of labeled *α*-SMA fluorescence to the mean response in the control group with [A]=0. The change in *α*-SMA expression above the mean control response (*E*–1) was fit by a Hill equation using a least-squares regression, yielding 3 parameter estimates: the maximal change in expression (Δ*E*_max_), the Hill coefficient (*n*), and the half-maximal effective concentration (EC_50_). Data is mean ± SD, N=3. One-way ANOVA conducted with Dunnett compared to 0 adiponectin control, *p<0.05.

### Adiponectin at concentration found in ACM only weakly induces myofibroblast conversion

We sought to determine if we could replicate the same level of *α*-SMA expression using ex-Adip at the same concentration native adiponectin is found in ACM (∼10 ng/mL). While ACM again induced significant and robust *α*-SMA expression, ex-Adip induced significantly less myofibroblast conversion (Fig. 5). Because ACM also contains ∼ 270 pg/mL of TGF-β1, we also tested whether there is a synergy between adiponectin and TGF-β1 by matching both adiponectin and TGF-β1 concentrations in ACM using ex-Adip and recombinant TGF-β1. This combination also failed to replicate the same level of myofibroblast conversion observed with ACM (Fig. 5). Thus, while ex-Adip is able to induce myofibroblast conversion, it fails to account for the full myofibroblast inducing potential of ACM. One potential explanation is that ex-Adip differs in form and/or lipid binding partners compared to the adiponectin in ACM. SDS-PAGE from the supplier shows several distinct bands, which suggests that ex-Adip contains a mix of different adiponectin forms and/or post-translational modifications.

**Figure 5.**
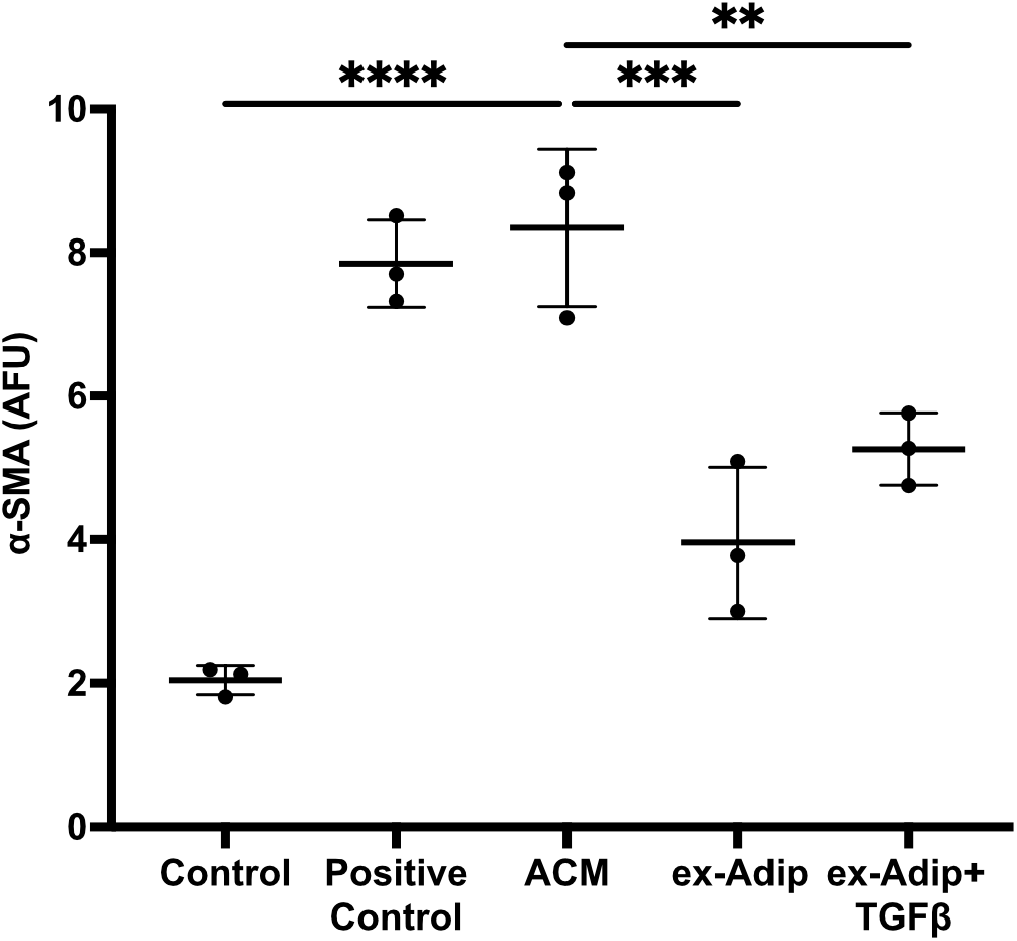
Exogenous adiponectin and TGF-β1 does not account for ACM mediated induction of myofibroblasts. Human adiponectin alone (ex-Adip) or adiponectin with TGF-β1 (ex-Adip+TGFβ) were added to 0.5% FBS DMEM in the same concentrations measured in ACM (∼10 ng/mL and 270 pg/mL, respectively) to determine if they synergistically contribute to myofibroblast conversion. Data is mean ± SD, N=3 One-way ANOVA with Dunnett’s multiple comparison compared to ACM, ****p<0.0001, ***P<0.001 **p<0.01.

### Depletion of native adiponectin from ACM significantly reduces myofibroblast conversion

Since ex-Adip could be different from the adiponectin in ACM, we next asked if depletion of native adiponectin from ACM would result in a loss of the myofibroblast converting potential. Using anti-adiponectin antibody coated beads, we depleted adiponectin from our ACM and repeated the *α*-SMA assay. We found that depletion of native adiponectin from the ACM resulted in a substantial and significant decline in fibroblast *α*-SMA expression (Fig. 6), which suggests that native adiponectin is necessary for ACM to efficiently induce myofibroblast conversion. To test if a second factor in ACM synergizes with adiponectin, we also spiked ex-Adip into the adiponectin-depleted ACM but it was unable to restore *α*-SMA expression back to levels seen with ACM. This finding suggests that the attributes of native adiponectin (*e*.*g*., it’s form or binding partner) are vital for mediating the conversion of fibroblasts to myofibroblasts. The different forms of adiponectin (*e*.*g*., globular, trimers, hexamers, multimers) are thought to each have distinct functions, including as carriers for other molecules, such as lipids ^11^. Furthermore, in addition to the total amounts of adiponectin, the distribution of the forms of adiponectin also has been attributed to different downstream biological functions^15^.

**Figure 6.**
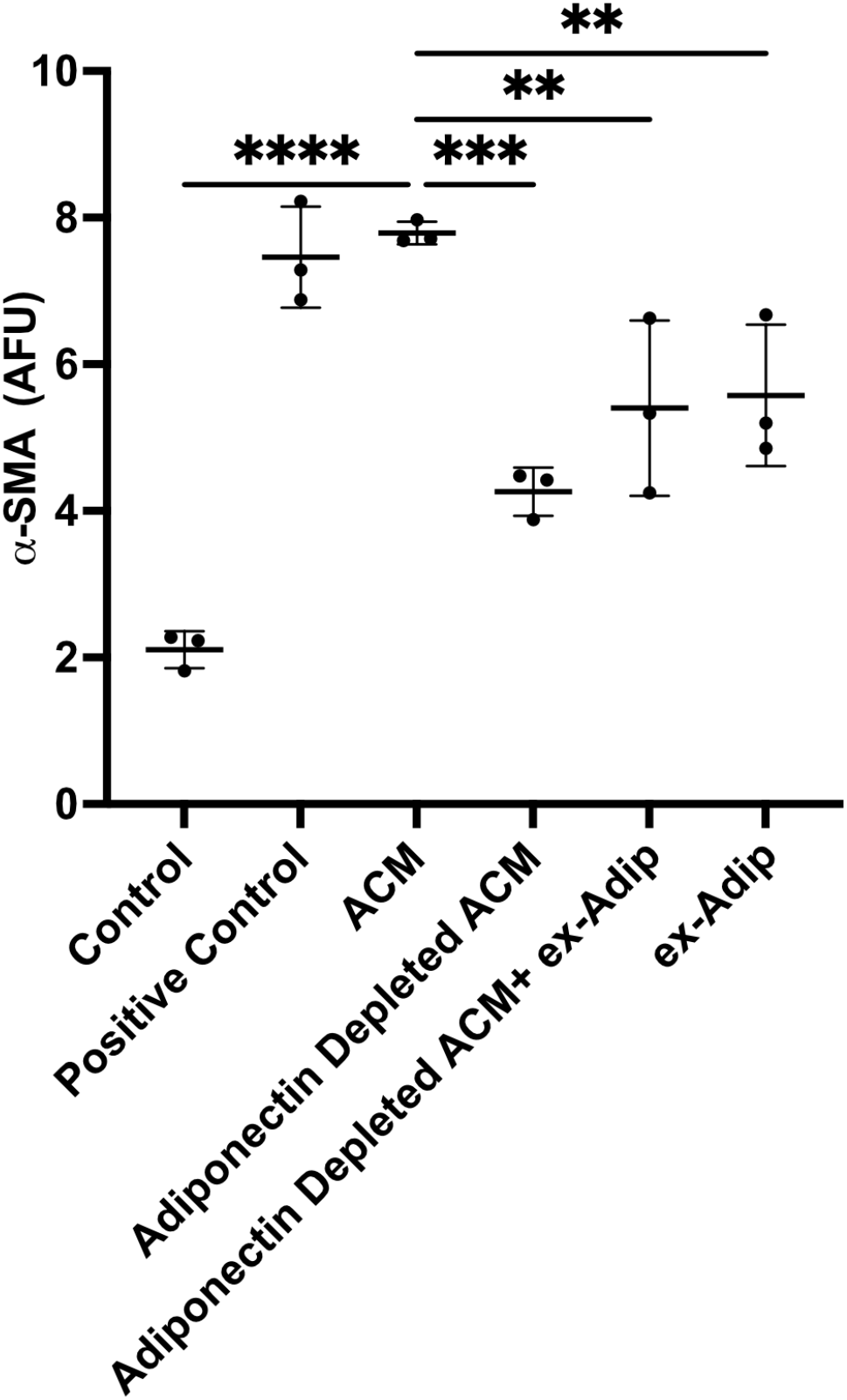
Depletion of native adiponectin from ACM substantially reduces ACM’s ability to induce myofibroblast conversion. ACM was depleted of adiponectin (Adiponectin Depleted ACM) using magnetic beads coated with anti-adiponectin antibodies and then applied to fibroblasts. Human adiponectin (ex-Adip) was added in the same concentrations measured in ACM (∼10 ng/mL) to determine if the function of native adiponectin could be replaced with exogenous adiponectin. Data is mean ± SD, N=3 One-way ANOVA with Dunnett’s multiple comparison compared to ACM, ****p<0.0001, ***P<0.001 **p<0.01.

## Discussion

Adiponectin is most commonly thought of as a hormone that regulates insulin resistance, but it is also associated with anti-inflammatory properties, as plasma levels are inversely correlated with many obesity-related inflammatory diseases^16^. In terms of direct anti-inflammatory effects on fibroblasts, adiponectin (and adiponectin agonists) has been shown to inhibit myofibroblast conversion^17,18^, mitigate collagen gene expression in the presence of TGF-β2^19^, and reduce ECM protein synthesis^20^, most likely through inhibition of the NF-κB pathway through AMP activated protein kinase (AMPK) signaling^16^.

In contrast, several clinical studies have found high adiponectin levels in pro-inflammatory conditions such as rheumatoid arthritis and joint disease^21-23^. Some *in vitro* studies also support a link between adiponectin and cell activities associated with inflammation. Adiponectin can induce *α*-SMA expression in monocytes^24^ and increase collagen production and hyaluronic acid (HA) synthesis in neonatal dermal fibroblasts^25^. There is also evidence that the form of adiponectin can impact fibroblast activity. Akazawa *et al*. found that HA synthesis in human dermal fibroblasts increased in a dose-dependent manner with exposure to recombinant oligomeric adiponectin but not to globular adiponectin^26^. Conflicting evidence on the effects of adiponectin on fibroblast/myofibroblast conversion might be due to a lack of exact details on the source and form of adiponectin used. For example, recombinant adiponectin can be made in either bacterial or mammalian expression systems, which can impact post-translational modifications known to impact adiponectin’s bioactivity^27^.

The apparent contradictions in adiponectin’s activity could also be due to the complexities of the fibroblasts themselves, as: (1) fibroblasts have behaviors specific to their tissue of origin^28^; (2) myofibroblasts within a tissue can originate from multiple cell sources^29^; and (3) myofibroblasts within a healing wound can be subdivided into several transcriptionally distinct subpopulations^3,4,30^. In light of these points, a more nuanced view of myofibroblasts is beginning to emerge. For example, myofibroblasts that originate from dermal adipocytes are enriched for genes associated with various ECM proteins, but not for genes associated with collagen maturation and cross-linking ^3^. These results suggest that healing outcomes could be dependent on which myofibroblast subpopulations are present in the healing wound. Therefore, an understanding of how those different subpopulations emerge and what their specific roles are could be beneficial for improving healing and tissue regeneration.

It is also worth noting that plastic and reconstructive surgeons have reported that fat grafting, which contains adipocyte stem cells, pre-adipocytes, and adipocytes, can reduce existent scar tissue through mechanisms that remain unknown ^31,32^. One possible mode of action is that these cells secrete factors that select for myofibroblast subpopulations that are more conducive to scar resolution. If this proposition holds true, it will be interesting to know if the adipocyte-mediated myofibroblast conversion we have observed, which occurs through a TGF-β1 independent mechanism, leads to a myofibroblast subpopulation that is either pro-scar or pro-regeneration. Such information will be important for optimizing current fat grafting techniques and for developing new therapeutic strategies.

In conclusion, physiochemical characterization of ACM revealed that the factor responsible for myofibroblast conversion is retained in the 30-100 kDa fraction and loses activity with heat treatment and lipid depletion. Significant differences in the secretion profile of common adipokines were found between ACM and PCM, but of the adipokines measured, only adiponectin has physiochemical properties consistent with our findings. For this reason, we investigated whether adding adiponectin at levels comparable to those measured in ACM could induce myofibroblast conversion. We found that fibroblasts are responsive to adiponectin in a dose dependent manner but at much higher concentrations than those measured in ACM. We also found that depleting native adiponectin from ACM removes its ability to induce myofibroblast conversion, suggesting adiponectin in its native form is important for its activity on fibroblasts.

## Materials and Methods

### Pre-Adipocyte and Adipocyte Spheroid Culture, Maintenance, and Conditioned Medium Collection

Pre-adipocyte spheroids were generated using a standard hanging drop cell culture as previously described^8^. Spheroids (each containing 20,000 cells) were then transferred to ultra-low adherent 24-well plates (Corning CLS3473), with 5 spheroids per well. Pre-adipocyte spheroids were maintained in pre-adipocyte growth medium (PGM-2, Lonza) containing 10% fetal bovine serum (FBS), 0.01% gentamycin sulfate /amphotericin B, and 200 μM L-glutamine for 10 days. Adipocyte spheroids were formed by culturing spheroids in pre-adipocyte differentiation medium (PDM-2, Lonza) containing 1% insulin, 0.1% 3-isobutyl-1-methylxanthine (IBMX), 0.1% dexamethasone, and 0.2% indomethacin for 10 days. After 10 days, medium was removed, the spheroids were rinsed gently with 1X PBS, then 0.75 mL of Dulbecco’s Modified Eagle Medium (DMEM, Gibco) containing 0.5% fetal bovine serum (FBS, Gibco 26140079), 1% penicillin/streptomycin (PS, Gibco 15140-122) and 0.1% amphotericin B (AB, Gibco 15290-018) was added to each well. After two days, this medium, termed either pre-adipocyte or adipocyte conditioned medium (PCM or ACM, respectively) was collected and stored at -80° C.

### LegendPlex Adipokine Panel

PCM and ACM were screened with a multiplex adipokine assay in order to measure levels of common cytokines, hormones, and other factors commonly secreted by adipose tissues. Collected medium was processed and run in triplicate via flow cytometry as described by the manufacturer (Biolegend LegendPlex Human Adipokine Panel, 741062).

### Protein Fractionation of Conditioned Medium

To fractionate ACM into pools of medium with different molecular weight cutoffs, a 100 kDa MWCO Vivaspin PES filter (Sartorius) was pre-rinsed with deionized water to ensure full saturation of the filter membrane. After removal of the water, ACM was added and centrifuged in a swing-bucket centrifuge for 8 minutes at 4,000 g per the recommendation by Sartorius. The column was inverted, re-centrifuged at 3,000 g for 2 minutes, and the concentrate was resuspended in 0.5% FBS DMEM to the original volume. To fractionate proteins in the 30-100 kDa molecular weight range, an Amicon 30 kDa MWCO (Milipore Sigma) filter was pre-rinsed as described before. The previously collected filtrate was added to the column and centrifuged at 1,500 g for 30 minutes. The filtrate was collected for additional fractionation steps. The concentrate was again resuspended in 0.5% FBS DMEM to the original volume. This process was repeated twice more with 10kDa and 5kDa MWCO Vivaspin PES filters (Sartorius), for a total of 5 fractions of ACM.

### Lipid Depletion & Heat Inactivation

To heat inactivate proteins in the medium, ACM was heated for 60 minutes at 85°C with a benchtop hotplate. For lipid depletion, ACM samples were treated with Cleanascite HC to remove lipid as described by Castro *et al*.^33^. 0.5 mL of Cleanascite HC was added to Eppendorf tubes and centrifuged at 1,000 g for 20 minutes to pellet the beads. The supernatant was removed, and the pellet was resuspended in ACM using a benchtop vortex. Samples were then incubated at 4°C overnight on a rocker plate. The next day, samples were centrifuged at 1,000 g for 45 minutes to pellet the lipid-bound beads. The lipid depleted ACM was decanted into a separate Eppendorf tube and filtered through a 0.45 μm pore size filter to remove residual polymer particles.

### Quantification of *α*-SMA expression

28-year-old human dermal fibroblasts from breast surgical discard (University of Iowa Tissue Procurement Core) between passages 3-5 were plated at 100,000 cells/cm^2^ in black walled 96 well plates containing 150 μL of DMEM with 10% FBS, 1% penicillin/streptomycin, and 0.1% amphotericin B. Fibroblasts were left to attach for 6-8 hours before removing the medium, rinsing with 1x PBS, and replacing with the various treatment groups in triplicate. The treatment groups included ACM and positive control (0.5% FBS in DMEM plus 1 ng/mL TGF-β1 and 50 μg/mL ascorbic acid) subjected to molecular weight fractionation in the ranges of >100kDa, 30-100kDa, 10-30kDa, 5-10kDa and <5kDa, and lipid depleted and heat inactivated. Fibroblasts were maintained in a humidified 95%/5% air/CO_2_ incubator at 37 °C for 48 hours. After treatment, samples were rinsed three times with 1X PBS, fixed and permeabilized with 4% paraformaldehyde and with 0.1% TritonX-100, respectively. Each well was blocked with 5% BSA-tween solution and incubated with a primary mouse anti-human *α*-SMA antibody (ab7817, Abcam, Cambridge, UK) solution at a 1:100 dilution overnight at 4°C. Samples were blocked again and incubated with 2-3 drops of an anti-mouse poly-HRP-conjugated secondary antibody from the Invitrogen Tyramide SuperBoost secondary staining kit (ThermoFisher, B40912). Samples were rinsed three times with 1X PBS and the tyramide working solution conjugated to AlexaFluor 488 was added and allowed to react for 20 minutes. Stop solution was then added followed by another three times rinse with 1X PBS. Cell nuclei were stained using Hoescht 33342 (Invitrogen, H3570) prior to imaging. The percentage of *α*-SMA positive cells was quantified using on a VarioSkan Lux plate reader. Measurements were made at an excitation/emission of 496/524 nm with each well receiving 29 measurements, each for 1000 ms.

### Addition of Exogeneous Human Adiponectin

To investigate the effects of exogeneous adiponectin on dermal fibroblasts, we reconstituted adiponectin from pooled human serum (BioVendor RD16202350) in sterile deionized water to a stock concentration was 0.1 mg/mL. Adiponectin was then added to low serum medium to produce final concentrations ranging from 6.125 ng/mL to 10 μg/mL for *α*-SMA quantification.

### Adiponectin Depletion from ACM

Using a DynaBeads co-immunoprecipitation kit (Invitrogen 14321D), 2 mg of magnetic beads were vortexed with C1 buffer for 30 seconds. The beads were collected with a magnet and conjugated with 12 μg of adiponectin antibody (ThermoFisher 19F1) in accord with the Invitrogen protocol. Conjugation was followed by a series of wash and collection steps using the supplied buffers. The final 10 mg/mL suspension of antibody-coupled beads was washed with 0.1% BSA in 1X PBS, then resuspended in 3 mL ACM. The solution was incubated at 4 °C with constant agitation for 10-30 minutes. The beads were collected with a magnet and the supernatant (adiponectin depleted ACM) was removed into a separate tube. Adiponectin depletion was confirmed using an ELISA (<0.3 ng/mL, Biolegend Adiponectin ELISA Max, 442304) and then the adiponectin depleted ACM was used in the *α*-SMA fluorescence assay to treat fibroblasts. Adiponectin from human serum was also added back to a portion of the adiponectin depleted ACM (10 ng/mL).

### Statistics

All statistical analysis was conducted in GraphPad Prism version 9 (GraphPad, San Diego, CA). Replicates from independent experiments were used to determine individual experiment means, which were then analyzed for significance (p<0.05) using either one-way or two-way analysis of variance (ANOVA) followed by either Dunnett’s multi-comparison tests or Tukey *post hoc* tests.

## Author Contributions

Mariam El-Hattab: Conceptualization, Methodology, Formal analysis, Investigation, Data

Curation, Writing – Original Draft

Noah Sinclair: Formal Analysis, Investigation, Data Curation

Jesse Liszewski: Methodology, Formal analysis, Investigation

Michael Schrodt: Methodology, Formal analysis, Investigation

Jacob Herrmann: Methodology, Formal analysis, Investigation

Aloysius Klingelhutz: Conceptualization, Resources, Funding Acquisition

Ed Sander: Conceptualization, Methodology, Formal analysis, Resources, Data Curation,

Writing – Review & Editing, Visualization, Supervision, Funding Acquisition.

James Ankrum: Conceptualization, Methodology, Formal analysis, Resources, Data Curation,

Writing – Review & Editing, Visualization, Supervision, Funding Acquisition.

## Acknowledgments

This study was primarily supported by a Pilot Grant awarded by the Iowa Technology Institute (EAS, JAA). Additional support from NIH P42 ES013661 (JAA, AJK), NIH R21 AR075137 (EAS), and a Diabetes Action Research and Education Grant (JAA). Adipose was supplied by the Tissue Procurement Core at the University of Iowa which is supported by an award from NIH (NCI award number P30CA086862) and the University of Iowa Carver College of Medicine.

